# Temporal dynamics of honeybee learning and decision-making are revealed by a novel automated PER system

**DOI:** 10.1101/2023.06.10.544458

**Authors:** Heather Strelevitz, Ettore Tiraboschi, Albrecht Haase

## Abstract

The proboscis extension response (PER) has been widely used for decades to evaluate honeybees’ (*Apis mellifera)* learning and memory abilities. This classical conditioning paradigm is traditionally administered manually, and produces a binary score for each subject depending on the presence or absence of the proboscis extension in response to a stimulus - typically an odor which has been associated with a sucrose reward - to classify whether or not the bee has learned the association. Here we present a fully automated PER system which delivers stimuli in a more controlled manner, and thus standardizes the protocol within and between labs; further, the AI-facilitated behavioral scoring reduces human error and allows us to extract a richer meaning from the outcome. The automated frame-by-frame assessment goes beyond the binary classification of “learned” or “not learned”, expanding the possibilities for many other measures. Using this method, we investigate the real-time decision-making processes of honeybees faced with difficult learning tasks. When posed with a quantitative (rather than qualitative, as in the case of different odors) PER association, honeybees show a pattern of rapid generalization to both the rewarded and non-rewarded stimuli, followed by a slowly acquired discrimination between the two. Our work lays the foundation for deeper exploration of the honeybee cognitive processes when posed with complex learning challenges.

## Introduction

Learning and decision-making are core features which allow an animal to survive in a stimulus-rich and dynamic environment. The cognitive capacities necessary to support these abilities are present not only in what we consider to be higher-order mammals, but also in organisms as far down the phylogenetic tree as the tiny invertebrate *Caenorhabditis elegans*[1]. The honeybee (*Apis mellifera*) is a particularly suitable model for the study of learning and decision-making considering their skill of foraging: a high-stakes, fast-paced challenge which the average adult bee performs with remarkable success. In a field of various floral signals, a honeybee must choose the targets with the highest chance of reward, and do this as quickly as possible. Underlying the behavior are complex interactions between sensory perception, sensory processing, memory, and motor systems - not to mention a continuous learning process. The mechanisms by which the honeybee efficiently optimizes this task has intrigued researchers for decades.

Here, we aimed to investigate these topics with new improvements to an old method. Since its introduction in 1961 [2], the proboscis extension response (PER) has been thoroughly characterized [3] and used with great success to probe the bases of insect learning through a classical conditioning paradigm which is both simple and versatile [4]. Within just a few trials, a harnessed honeybee will learn the association between a conditioned stimulus (CS) and an appetitive unconditioned stimulus (US), usually sucrose. Typically this experiment requires the researcher to manually provide all stimuli: the sucrose solution would be supplied to the bee from a syringe, and the odor would be delivered from syringes containing filter paper imbued with the odorant. This would be followed by recording the observed dichotomous behavioral response, which quantifies whether or not the bee extends the proboscis to the CS before the presentation of the US [5]. Experimenting with PER takes advantage of the bees’ natural reflex to extend the proboscis when the antennae, tarsi, or mandibles come into contact with sucrose [4], which in healthy bees is a robust and reliable phenomenon. This protocol allows for the investigation of both sensory perception and the abilities of learning and memory, and is widely used due to its ease of implementation and low-cost equipment.

However, there are a variety of protocol details which are known to cause variation in the PER acquisition, and are difficult to control manually: stimulus intensity, air flow, and timing of stimulus presentation, to name a few [6]. Variations among these factors may reduce the validity of comparing data between different labs, or even within the same lab between experiments run by different researchers.

Further, the binary classification of behavioral output – often necessary for a manual experiment – is only one piece of the information available, and thus limits the scope of our biological comprehension.

In recent years, several efforts have been made to implement automated tracking of honey bee head parts to facilitate experiments and to obtain quantitative information on the bees’ responses to stimuli. Many of the efforts concentrated on tracking antennal motion. A first attempt used paint-marking of antenna tips, which allowed their straightforward identification on the camera images [7]. This showed how odor conditioning influenced the antennal motion pattern. Another study with markerless tracking of the antennae and proboscis tip used the large contrast between the tip and the background, which was detected via an algorithm that followed pixel-by-pixel average color changes [8]. Currently the gold standard in animal tracking is the deep learning python toolbox DeepLabCut [9]. The strategy here is transfer learning from a very deep neural network pre-trained on huge object recognition datasets. Antennal tracking experiments have begun to take advantage of this tool [10], again analyzing antennal motion in response to odor stimuli.

Efforts towards fully automatized conditioning have also been made before. One approach performed automatic analysis paired with manual conditioning using bayesian-based algorithms to identify the motion of antennae, mandibles, and proboscis [11]. Another study implemented a fully automated approach for aversive conditioning within a walking arena, where the walking motion of the bees could be continuously monitored and electric shocks could be provided via the walking surface [12]. In the presented work we addressed both problems: the complete automation of the administration of both stimuli (CS and US) in a fully customizable sequence to multiple subjects; and real-time response detection supported by machine learning classification of the proboscis extension. The device was tested in a series of experiments that reveal new insights into the temporal dynamics of the behavioral responses.

## Results

### Classical PER conditioning

The first experiments were classic olfactory conditioning paradigms, where one odor was rewarded with a sucrose solution and another was not. In both cases, the movements and timing of the machine and its stimuli were the same; the only difference was the rotation of the feeder from the left (containing sucrose solution), or from the right (empty). Both of the odors had a neutral valence for honeybees, as evidenced by similar results regardless of which one was rewarded. The paired and unpaired trials were presented in a pseudo-random order over ten trials, such that each subject receives trial 1 before moving on to trial 2, resulting in an 8-minute inter-trial interval for a single bee (Figure 1a).

**Figure 1:**
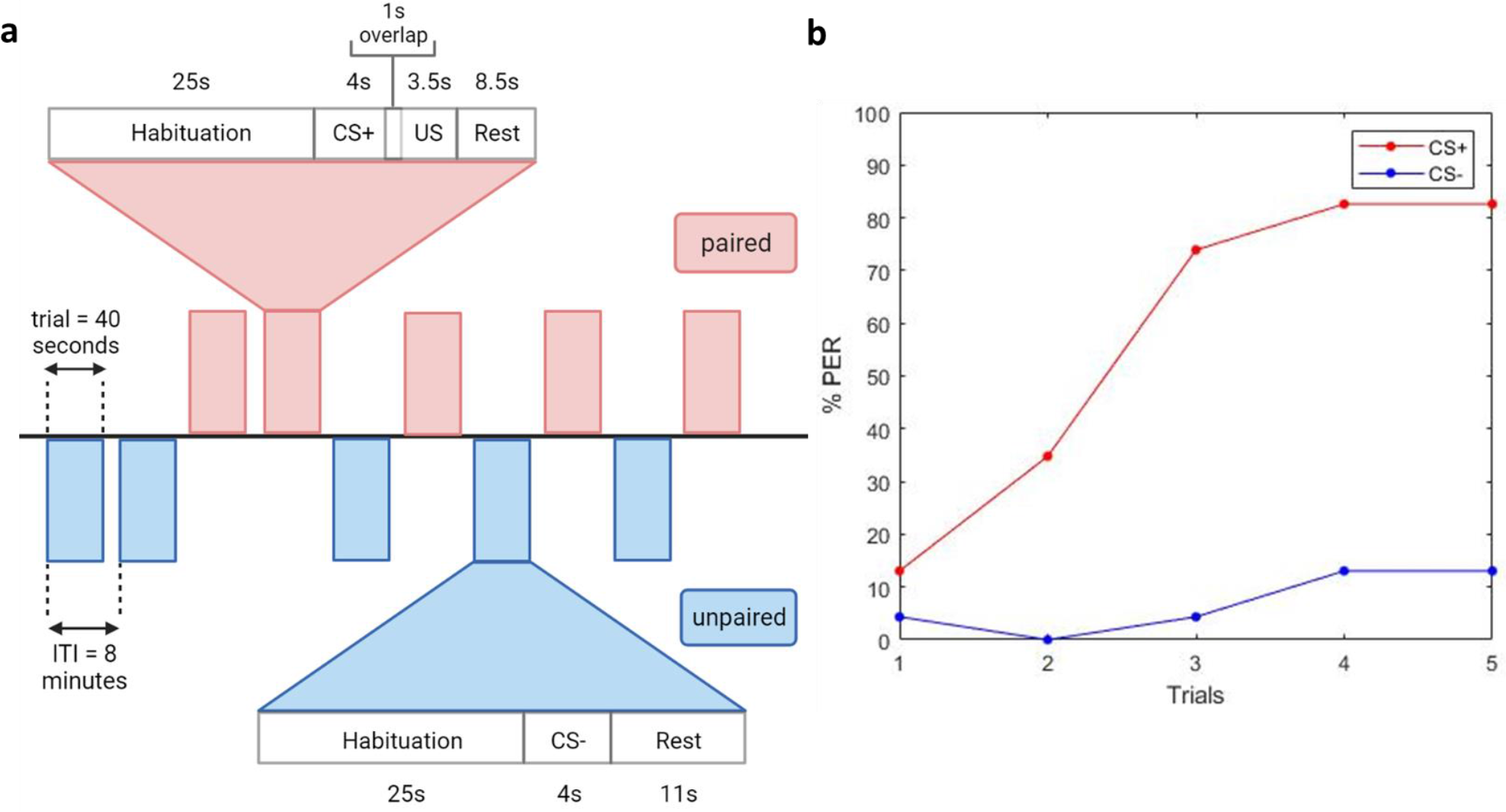
A proof-of-concept demonstration of an olfactory conditioning experiment with the automated system. **a)** A protocol diagram representing one complete experiment with 10 pseudo-randomized trials. The timelines are depicted for one trial of each of the paired and unpaired paradigms. One trial contains the time between the bee moving into the test position and moving out of it. The inter-trial interval (ITI) is the amount of time between one trial beginning to the next trial beginning, for one individual bee. This is also indicative of one complete rotation of the wheel holding the 12 bees. **b)** Classic PER response curve over trials. *n* = 24. 1-hexanol (red) was rewarded (CS+), 1-nonanol (blue) was not (CS-). Each was diluted 1:100 in mineral oil.

Evidently, our automated system achieves the same success as previous manual or semi-automated setups. By the end of three rewarded trials, the majority of subjects (74%) had learned the association between the odor and the sucrose, as quantified by the extension of the proboscis to the odor presentation prior to the sucrose delivery. This was calculated using the CNN classifier: at every camera frame the classifier outputs a probability that the proboscis is extended, and we set a threshold (0.8) over which we categorize the frame as having the proboscis extended. We removed from the analysis any subjects which did not extend the proboscis to the antennal sucrose stimulation, since if they have lost their innate PER, we cannot use their PER as a behavioral metric for learning. We also removed subjects which already had the proboscis extended for greater than half of the second prior to the onset of the CS, to avoid classifying as “learned” those subjects which already had the proboscis extended by chance when the CS began. With the remaining subjects, within the time window of the CS which did not overlap with the US, we classified a subject as having learned when the proboscis was extended for one quarter of a second or longer.

By the last paired trial, the PER was 83%. Meanwhile, the unpaired odor did not elicit the same learned association response, never reaching a higher percentage than 13% (Figure 1b). Statistical analysis shows a significant main effect between groups (*F*(1,46) = 35.9, *p* = 3×10-7). Simple within-subject effects show learning in the CS+ group (*F*(4,92) = 22.2, *p* = 7.3×10-15) but not in the CS-group (*F*(4,92) = 0.68, *p* = 0.61). This group dependence of learning is also reflected in a significant interaction of group × trial (*F*(4,184) = 19.3, *p* = 2.7×10-13). During the first trial, the difference between groups is not significant (*F*(1,130) = 0.015, *p* = 0.91), but this difference becomes significant already at the 2nd trial (*F*(1,130) = 4.93, *p* = 0.036) (see electronic supplementary material, Table S9).

### Mechanosensory stimuli

To further explore the stimulus possibilities with our setup, we ran a PER experiment using stimuli which were not two different odors but rather two different magnitudes of air flux (without odor). We consider this task to be more challenging because rather than being distinguishable by chemical compounds, as in the case of odors, different air fluxes are the same quality of stimulus presented in different quantities. The protocol was the same as for the experiment in Figure 2, except for the type of stimulus and the number of trials (which was increased from 10 to 32 to allow more time for learning). Here the lower flux was rewarded (CS+) and the higher flux was not rewarded (CS-). Notably, the classic PER learning curve (Figure 3a) is similar to a simple olfactory PER experiment: 83% of honeybees succeeded by the third CS+ trial, peaking at 92% on trials 5, 6, 10, and 14; meanwhile the response to the CS-was never higher than 17%.

**Figure 2:**
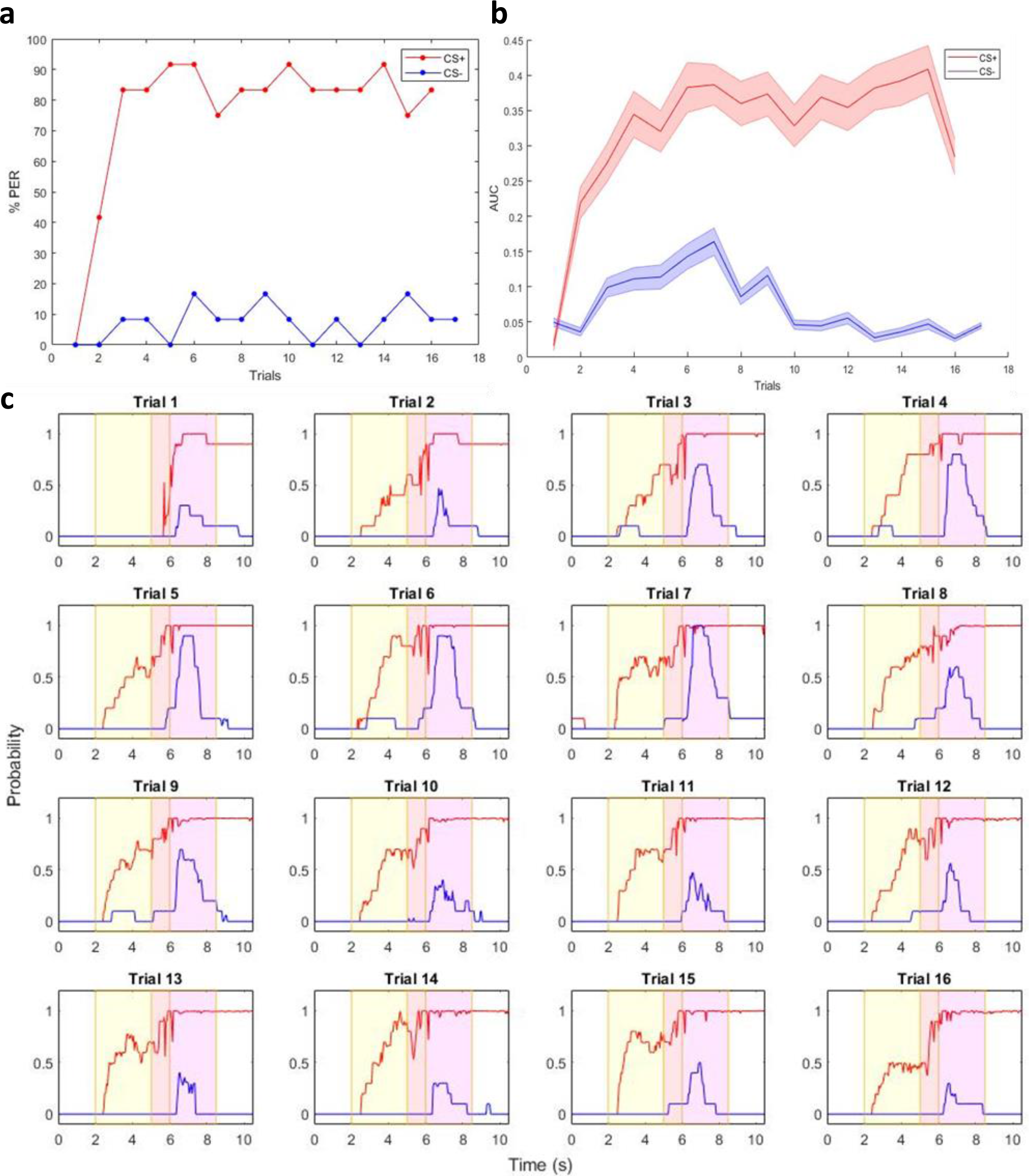
Honeybees successfully perform PER with two different (odorless) air fluxes, where the lower flux is rewarded. **a)** Classic PER response curve over trials. The low air flux stimulus (1.25 m/s), in red, was rewarded (CS+). The high air flux stimulus (5 m/s), in blue, was not rewarded (CS-). **b)** Area under the probability curves for CS+ and CS-stimuli for each trial. This is a measure for the probability over time, as given by the classifier, that a bee has the proboscis extended. **c)** A trial-by-trial view of the results in (b), where the probability of the proboscis extension is plotted over time in seconds. Results are averaged across all bees (*n* = 12). The yellow panel indicates the time duration of the conditioned stimulus (odor) and the blue panel is the time duration of the unconditioned stimulus (25% sucrose solution). There is a 1 second overlap.

**Figure 3:**
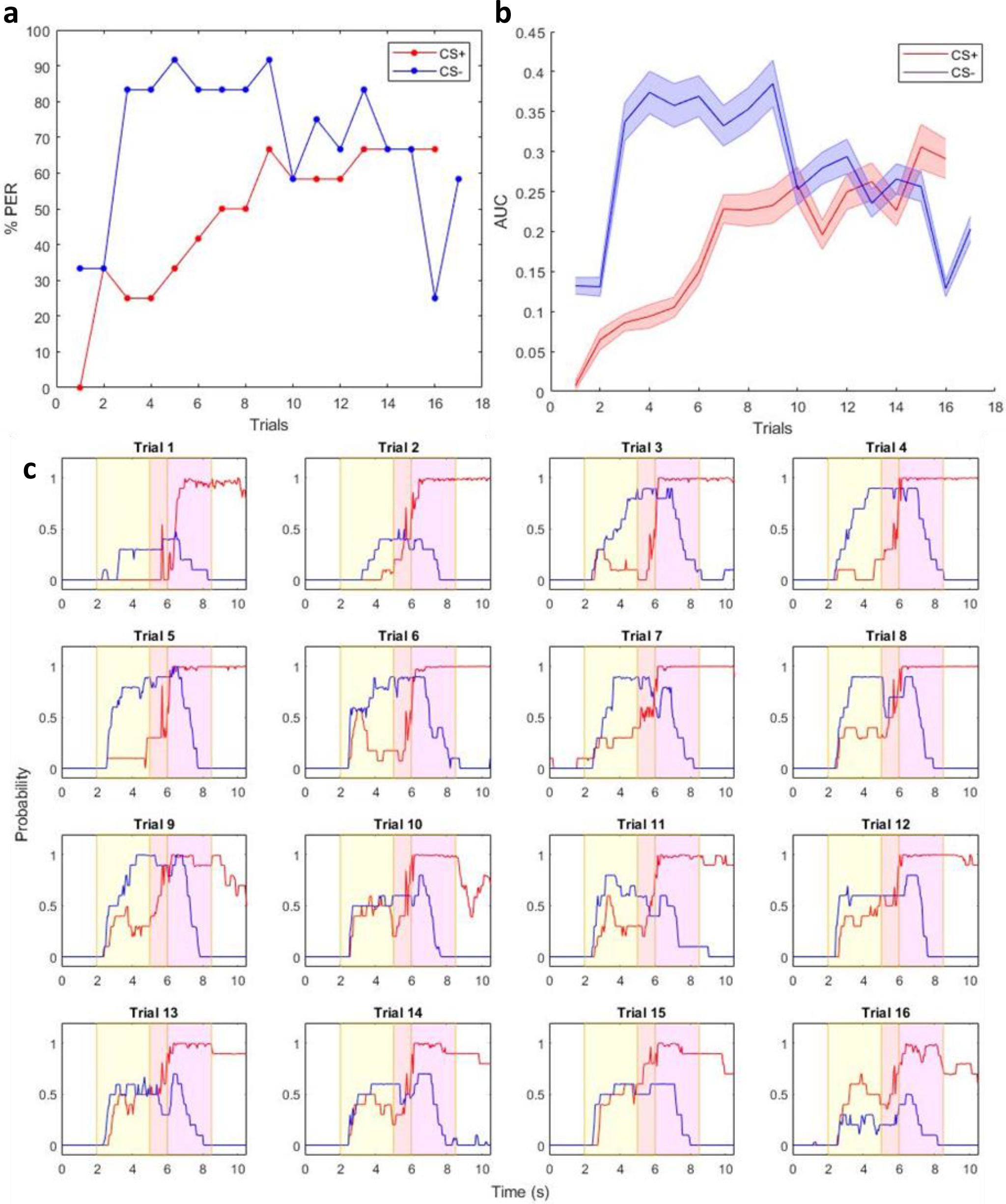
Honeybees are less successful with two different (odorless) air fluxes, where the higher flux is rewarded. **a)** Classic PER response curve over trials. The high air flux stimulus (5 m/s), in red, is rewarded (CS+). The low air flux stimulus (1.25 m/s), in blue, is non-rewarded (CS-). **b)** Area under the probability curves for CS+ and CS-stimuli during each trial. This is a measure for the probability over time, as given by the classifier, that a bee has the proboscis extended. **c)** A trial-by-trial view of the results in (b), where the probability of the proboscis extension is plotted over time in seconds. Results are averaged across all bees (*n* = 12). The yellow panel indicates the time duration of the conditioned stimulus (odor) and the blue panel is the time duration of the unconditioned stimulus (25% sucrose solution). There is a 1 second overlap.

In addition to the traditional categorical PER scoring, our setup allows for analysis of the response behavior in more detail. The probability of proboscis extension is available at every camera frame. The average time curves across bees are shown in Fig. 2c, where the CS period is shadowed yellow and the sucrose reward period (provided only to the CS+ group) is shadowed magenta. A quantitative analysis of the time-resolved process before and during proboscis extension would not be possible by manual observation.

The time averaging over single trial periods provides a mean area under the probability curve (Fig. 2b). For the CS+ group, this area is measured during the first 3s where only the CS+ stimulus (the odor) is provided since the proboscis extension during the sucrose administration is very stereotyped and therefore does not provide any further information. However, the CS-group shows important features also after the 3s, so that the area is analyzed across the whole 6s. To render the integrated probabilities comparable they are divided by time.

Looking at this temporally resolved behavior, learning in the CS+ group is revealed by a trial-by-trial reduction in reaction time to the CS+ stimulus, as well as an increased probability amplitude signaling greater proboscis extension overall (Fig. 2c, red curves). The integration over trial periods shows that a full consolidation of a stereotyped response requires 7 learning trials (Fig. 2b, red curve), which is not obvious from the classical learning curve (Fig. 2a, red curve). The CS-group shows hardly any response during the CS-stimulus, but after the stimulus is switched off, a curious tendency of proboscis extension is noticeable (Fig. 2c, blue curves, magenta region). This effect increases, clearly visible after averaging over the trial periods, until trial 7, but then decreases again to its initial level after 13 trials (Fig. 2b, blue curve).

The statistical analysis of the proboscis extension probability (per time) shows a significant main effect between groups (*F*(1,22) = 12.0, *p* = 0.0023). Simple within-subject effects are significant in the CS+ group (*F*(15,165) = 2.88, *p* = 4.8×10-4) and, different from the odor conditioning experiment, also in the CS-group (*F*(15,165) = 3.96, *p* = 4.5×10-6). The trial-dependent changes still depend on the group, manifested in a significant interaction group × trial (*F*(15,333) = 2.39, *p* = 0.0028). Simple between-subject effects become significant only from trial 7 (*F*(1,85) = 5.30, *p* = 0.038) (see electronic supplementary material, Table S9).

### Aversive mechanosensory stimuli

In a third experiment, we inverted the rewarded and unrewarded stimuli. Now the high air flux is associated with the sucrose reward, while the low flux is not. Both analysis methods, the classical binary evaluation of the proboscis extension (Fig.3a) as well as the PER extension probability curve (Fig.3b) show a similar picture. The unrewarded low flux (CS-) starts eliciting proboscis extension at a rapidly increasing rate arriving at 83% by trial 3. Compared to that the rewarded high flux (CS+) shows a much slower increase arriving at 25% by trial 3. However, over time the CS-curve steadily decreases while the CS+ curve increases, until after 16 trials the situation has inverted to 25% for CS- and 67% for CS+. The trial-by-trial temporal analyses (Fig.3c) show that after 3 trials the PER response onset time is almost identical for both stimuli and stays constant for the rest of the experiment. Only the PER probability amplitude continues to change over the trials: it starts to increase for the CS+ group from trial 6, and reduces for the CS-group from trial 9 until the areas under these curves have inverted at trial 16.

Statistical analysis of the proboscis extension probabilities across subjects (Fig. 3b) revealed a significant main effect between groups (*F*(1,22) = 5.38, *p* = 0.031), although smaller than for the previous experiment where the low-flux was the CS+. Simple within-subject effects are significant in the CS+ group (*F*(15,165) = 3.13, *p* = 1.7×10-4) and in the CS-group (*F*(15,165) = 3.32, *p* = 7.2×10-5). The trial-dependent changes again depend on the group: the interaction group × trial is significant (*F*(15,333) = 3.17, *p* = 7.1×10-5). Simple between-subject effects become significant from trial 3 (*F*(1,85) = 8.89, *p* = 0.02) (see electronic supplementary material, Table S9). The effect disappears again after trial 6, with one exception at trial 9 (*F*(1,85) = 7.84, *p* = 0.02).

## Discussion

The technical advantages offered by our automation are numerous. The delivery of stimuli can be precisely controlled both within and between subjects; an increased variety of stimuli is possible; and the code is easily customizable for individual needs. Furthermore, the internal validity becomes unquestionable when the entire protocol is automated and its parameters electronically saved, rather than relying on human performance and memory for administering the experiment and recording its data. Not only do these improvements allow for more easily standardized techniques - and therefore reliable comparison within and between labs - but they also reduce human error and bias during data analysis. Further, the precision of the control over the stimuli is crucial for maximizing the potential of the protocol. Of course there is also the added bonus of saving researchers’ time: once the protocol code is running, the system requires no further manual input (except, for longer experiments, periodically refilling the feeder containing sucrose solution).

Video recordings of PER experiments have been used for data analysis for over two decades, yet were always limited by the labor-intensive nature of manual scoring [13, 14]; meanwhile, our real-time analysis by a neural network allows researchers access to instant results. Original efforts to track paint-marked body parts in honeybees were more efficient than manually labeling individual video frames, revealing the influence of odor conditioning on antennal motion [7]. However, physical labeling is a substantial effort, and has since been improved by markerless tracking analysis techniques. These have been used to show antennal responses to novel odors [8]and to obtain detailed data on proboscis extension and motion onset [11]. Yet these methods also required significant manual efforts since the tracking algorithms needed to be trained for each subject. Further, although these approaches provide more information by precisely tracking body parts, they encounter severe problems when these body parts overlap. One paper reported error rates of 0-22% in the PER classification for single subjects [11].

Meanwhile, our model reaches a classification accuracy of 99.7% within its training sets. Typically when analyzing a new experiment, we will randomly check some videos against the model’s output to ensure validity; if there are any misclassifications, we retrain the model on videos from that experiment. However, this happens less and less frequently as the model is trained on data from different experiments.

The current gold standard for markerless pose estimation, DeepLabCut, has recently been used for the first time to track honeybee antennae, revealing a correlation between odor valence and antennal motion patterns [10]. For these experiments, the network was trained on multiple body parts for every single video; however, this required very few frames (3-10) to be labeled for each video. The approach presented here needed a far larger training set for our convolutional neural network, a few hundred frames, but rather than individually labeled body parts it requires a binary classification of whether or not the proboscis is extended in a given frame. Further, thanks to the excellent reproducibility of the head position among subjects, this training is required once and then the trained model works for all subjects. So far, DeepLabCut has not been tested specifically on the proboscis, and it may be a less ideal tool for such a body part which is only visible some of the time.

One approach that reported fully automated conditioning [12] overcame both limitations: automatic analysis of data as well as the mechanized stimulus generation. It was also shown to work with optical stimuli to study color learning [12]. We provide complementary information by working with aversive conditioning and analyzing proboscis movement. Additionally, in Kirkerud et al’s setup, experiments must be initialized subject-by-subject while our system performs a fully automated assay on 12 subjects in a single experiment. Thus, in this work, we have begun to expand the limits of the PER method’s capabilities. With a fully automated system, experimental sessions are no longer limited by the stamina of the researcher, and so can be multiple hours long, allowing us to propose difficult tasks and observe the bees’ behavioral changes over many trials. During our mechanosensation experiments, increasing the number of trials provided valuable information which would have been incomplete during shorter experiments; and only on the last couple of trials did we begin to see signs that the bees were becoming satiated (as indicated by slightly decreased AUC during the period of sucrose delivery).

The universality of the setup was then tested with a different type of stimulus. An increasing number of studies suggest that the insect olfactory system might not be limited to the processing of odor information, but is also sensitive to certain mechanical stimuli. Structural as well as functional evidence has been found at the level of the antennal lobe [15–18] as well as in higher-order brain centers [19,20]. Thus mechanosensation was an interesting candidate for PER experiments, particularly since other sensory modalities such as vision have been previously used but with less success than olfaction [21].

In an initial experiment we presented two different speeds of air flux at a ratio of 1:4, with the lower air flux equal to 1.25 m/s and the higher air flux equal to 5 m/s. This ratio was chosen according to animal research regarding numerosity, in particular Weber’s Law, which states that the ease of discriminating between two magnitudes depends on their ratio [15]. A lower ratio of 1:3 was tested and proved to be too difficult, with a very low rate of successful discrimination (data not shown). Increasing the ratio requires either a decreased low flux, which is limited by the technical capabilities of the valves, or an increased high flux, which could begin to depart from ecological relevance and become aversive (as we speculate below, we could already have an aversive element to our high air flux).

The results show a classical learning curve similar to that of the odor learning experiment (Figure 2a). Interestingly, the success rate more closely mimics that of olfaction experiments rather than vision experiments, possibly due to the findings regarding mechanosensation processing in the antennal lobe as discussed above. Furthermore, the novel measure of the proboscis extension probability (Figure 2b), which can be considered a rating of the PER distinctiveness on a given frame, allows for the detection of minor responses that would normally be simply classified as “not extended”. This reveals slight responses to the CS-stimulus at the beginning of an experiment which disappear after a few trials, suggesting a certain degree of initial generalization between the CS+ and CS-stimuli. The incomplete extension suggests indecision or uncertainty during the response, which is remarkable because PER was previously considered a reflex that is strongly triggered whenever a learned stimulus is identified. Yet this proposition is in agreement with recent evidence indicating that mechanisms of attention may underlie associative learning in insects [16]. Our closer look at the behavioral process suggests that PER is a behavior containing large variability, which reflects the degree of certainty with which the decision was made regarding stimulus identity; such a measure is particularly useful during the learning process when the animal’s certainty is low. This would remain unnoticed if only full proboscis extensions were counted by applying a threshold to the PER probability (Figure 3a). The tendency to generalize between stimuli during PER conditioning was not found in the vast majority of previous studies; it is possible that such a phenomenon went undetected due to the lack of sub-threshold scoring. However, in the work of Bitterman et al. [3] a similar small and fast decaying response to the CS-is present in odor conditioning experiments.

The time-resolved monitoring of the process (Figure 3c) gives further insights into the decision-making process: most of the subtle responses to the high flux CS-stimulus appear immediately after it is switched off. At that point, for a short moment, the decreasing air flux velocity passes the value of the lower flux CS+. This might cause a reevaluation of the stimulus leading to these timid responses. This temporal resolution will allow for the investigation of interesting aspects of decision-making, such as speed-accuracy dependence [17], since we can now measure with high precision the onset of right and wrong responses. Moreover, the shape of the response curve may provide information regarding the coding mechanism in the neurons involved in the decision-making process. Experiments in vertebrates show complementary contributions of latency and intensity coding [18], while in insects both coding mechanisms could be identified so far only in the periphery of the olfactory system [19]. It will also allow for the study of the temporal evolution of decision-making for stimuli that require temporal integration for their identification, like oscillations [20]. For honeybees, this is particularly interesting to address the open questions of how the waggle dance is decoded in the brain [21] [22].

In the second mechanosensory experiment, the stimuli were switched such that the CS+ was the high flux and the CS-was the low flux. The resulting data were surprisingly different from the previous experiment; so much so, that it is possible the stimuli are not similar enough in valence to be considered comparable for this type of experiment. We suggest that the high air flux is not a neutral stimulus, but a mildly aversive one, due to the fact that honeybees take many more trials to learn its association with a reward (whereas in the same context honeybees are able to much more quickly learn the same thing for a low air flux). Similar results are seen for PER experiments using other stimuli which are known to have a biological valence, such as the sting alarm pheromone (SAP), where the main component of SAP (isoamyl acetate) impaired the bees’ learning of the sucrose association [23]. This effect may be because, in ecological settings, proboscis extension (typically for feeding or foraging behavior) and the presence of SAP should be mutually exclusive. Thus, the bees need to learn to extend the proboscis to a stimulus which would normally inhibit such an action. In other experiments, where the SAP component was directly used as CS+ stimulus, the learning success was again strongly reduced and an unusual generalization could be observed: after conditioning to isoamyl acetate, bees exhibited PER for several novel odor stimuli with very different chemical properties [24]. A similar scenario was observed in the experiments presented here.

While the bees were conditioned to the apparently aversive high air flux, they reacted after a few trials with PER in response to the unrewarded low flux stimulus. They seem to completely generalize between the two stimuli: while the CS+ response is reduced due to its intrinsic negative valence, the CS-response induces PER with a learning rate as high as if it was the rewarded stimulus. Additional trials reversed the situation: while PER responses to the CS+ increase with time, the responses to the CS-gradually decrease.

Interestingly, the opposite task - bees learning to withhold the proboscis despite being stimulated with sucrose - has already been successfully performed [25]; together, these works demonstrate that responses to aversive stimuli can not exclusively be studied via the sting extension response (SER) protocol, but useful insight can be obtained via the PER paradigm as well. Future studies of complex learning and decision-making processes with conflicting stimuli will benefit from the possibility of resolving the temporal dynamics of individual responses. This provides a tool that allows for experiments which test the limits of insect cognition [26]. For example, one could test whether emotion-like states can have an influence on decision-making not only at a scale that changes the final outcome, as shown in a previous study [27], but also on a more subtle level wherein only the dynamics of the process may be altered. Adding other distractive stimuli would allow for studies on selective attention [28]. Additionally, studies on brain lateralisation may gain from an automated analysis with high temporal resolution. PER conditioning has contributed substantially to the discovery and characterization of left-right asymmetry in the honey bee brain [29, 30], but these projects always required two experiments with one of the antennae inactivated by a silicon coverage. We propose a more direct approach of providing stimuli from either the left or the right side while both antennae remain active. In summary, our flexible automated setup will allow researchers to expand their experimental repertoire to ask more subtle, complex questions, and to peer more deeply into honeybees’ learning and decision-making processes.

## Methods

### Honeybee Maintenance and Handling

No legal regulations apply to experimental research with honeybees. Subjects were maintained in the outdoor colony of the University of Trento at their site in Rovereto, TN, Italy. Experiments were performed from July to October 2022. Forager bees (as confirmed by the presence of pollen on the hind legs of a bee returning to the hive) were individually collected and immobilized on ice to allow for the mounting procedure described below. The duration of the contact with ice was minimized, using only the time required for sufficient handling.

### Hardware

The device comprises several elements (Fig. S1):

- Revolver: a wheel where the bees are mounted
- Feeder: provides the consumable stimulus as well as a tactile stimulus to the antennae
- Camera: records the behavioral response of the bee
- Stimulus generator: a 2-channel olfactometer (this independent module can be replaced by arbitrary stimulus generators such as visual, olfactory, auditory, mechanical, etc.)
- Controller: an Arduino-based controller that drives all the motorized components and if necessary also the module for the stimuli (see schematic circuit)
- USB Nidaq I/O port: the port of communication between the PC running the main program, the arduino controller and the actuators (valves)

All the structural components for the revolver were 3D printed and/or machined based on the technical drawings provided (Fig. S6). We used PETG (glycolized polyester), and TPU (Thermoplastic Polyurethane) for 3D printing, and plexiglass for machining. A scheme of the full assembly is provided (Fig. S5).

All electronic components are listed in the electronic supplementary material, Table S3, and their wiring is shown in Fig. S2. The wired Arduino and USB NI DAQ ports shown in this figure correspond to those addressed in the provided software (https://github.com/NeurophysicsTrento/Automatized-PER).

### Rotor and feeder

The device is based on a rotating wheel, the revolver, where 12 bees are loaded along the circumference facing outward (Fig. S1b). The bees are mounted on a tubular bee holder (Fig. S7b) after immobilization on ice and by gently grasping the thorax using tweezers. Once mounted (Fig. S7c), the holder allows the bee to extend its proboscis and freely move its antennae while holding the bee fixed by means of a head holder (Fig. S7a). A piece of soft sponge is placed on the back of the bee to prevent any body rotation. The revolver positions a single bee in front of the feeder and the stimulus generator. After each trial, the motor spins the revolver to place the next bee into the experimental position. The correct positioning is achieved by a rotary encoder that determines when to interrupt the rotation.

To perform sucrose conditioning, a servo motor rotates the feeder (Fig. S1a) in front of the bee to apply the US stimulus. This consists of touching the antennae with a sucrose-soaked stick and providing the 25% sucrose solution for feeding (Fig. S1c). Both are required to trigger PER association.

The covered revolver forms isolated compartments for each bee with a funnel-like shape. In the case of olfactory conditioning, the CS odor stimuli are removed from each compartment by means of a suction tube attached below the revolver (Fig. S1a). During the trial, a camera (Fig. S4) records a video of the bee from above, which is later used for behavior classification and data extraction.

### Odor stimulation

Odor stimuli are delivered using a custom-made 2-channel olfactometer. Briefly, a fluxometer feeds air through glass vials containing either an odor diluted in mineral oil or only the solvent. A fast 3-way solenoid valve selects which channel is opened. By default, the solvent channel is continuously streamed through a nozzle aimed at the head of the bee in the experimental arena. The CS odor stimulus is then delivered by activating a solenoid valve. The trigger is sent from the main program through the NI DAQ board to the circuit that powers the valve (in this case, a Darlington array chip (ULN2003)) (Fig. S2). Be aware of the appropriate voltage required by the valve; in our case, it is 12V for activation of the solenoid and 5V for holding. Due to the short period of the stimulus, a VCC of 12V can be used throughout the whole stimulus.

### Mechanosensory stimulation

In order to control the air stream intensity we used a voltage-regulated valve (xxx), such that the air flux is controlled by varying the voltage. The necessary current for operating the valve is provided again through a channel of the ULN2003 chip. The gate of a channel on the ULN2003 must be connected to a PWM pin on the Arduino and the voltage-regulated valve must be connected to the ULN2003 as the odor valve (Fig S2). For the experiments presented here, a separate Arduino was used for controlling this valve simply to obtain higher modularity of the system. A calibration of the valve is necessary in order to define its working range, which is based on the input air pressure. In this case, the Arduino is driven by digital inputs received by the main program through the NI DAQ board. The flux is measured with a thermo-anemometer (Testo 405i, Testo).

### Software

In order to execute an experiment, the device requires 3 layers of software:

- Main program: written in Matlab (R2019b, Mathworks), it oversees the entire device functions according to the experimental protocols, as well as analyzing the resulting data.
- CNN model: a convolutional neural network that identifies the behavioral response of the recorded bee. It has been generated and trained using TensorFlow and Keras framework. The network is then imported into the Matlab environment for live classification of bee responses.
- Hardware controller code: written using the Arduino IDE, it is meant to control all the motors and actuators for stimulus controls (valves, solenoids, speakers, LEDs, etc.). It can be adapted for different experimental protocols since it controls the synchronization, timing, and type of stimuli.

All the codes are commented with user instructions and can be found at https://github.com/NeurophysicsTrento/Automatized-PER. In the case of the CNN model, the datasets used for the training, and the trained model itself, are provided along with the Matlab code used to manually create labeled datasets from recordings of PER experiments. The software functions are summarized as follows:

### Main code

The main code, written in Matlab, oversees the entire PER experiment. It initializes the USB NI DAQ port for communicating with the hardware; it initializes the camera and controls the video recording and analysis; it defines the exact protocol for the PER conditioning, such as the sequence and timing of all stimuli (PER_Protocol.m); and it saves the final dataset as a .mat file which contains all the parameters of the experiments, the recordings, and the classifications by the AI model. The main analysis script (analyse_PER.m) then uses this file to perform analyses and visualizations of the experimental data. Also, it includes functions to extract important parameters like the classical learning rate, and contains visualization tools to help summarize the outcome of the experiment and to highlight behavioral features. It then allows a revision of the experimental data before finalizing the analyses.

### Convolutional Neural Network

The CNN performs an automated classification of the behavioral responses of the bees. During the trial period, a video of the subject is recorded and automatically fed to an AI model for classification. The classification is based on the extension of the proboscis: the response is defined as “licking” if the end of the proboscis is extended beyond the mandibles of the bee, or “rest” otherwise. When the proboscis tip is in close proximity to the mandibles’ edge the classifier will be less accurate, since the manual classification of those frames will not always be scored in exactly the same way between different videos and experimenters. The output of the model is the probability that the proboscis is extended, a value between 0 and 1 that is assigned to each frame. The model is a convolutional neural network, generated using Tensorflow and the Keras framework, which takes inputs as frames of size 100×100 pixels. The provided colab notebook script (PER_CNN.ipynb) can be used to train and retrain a model. In general, the model requires a training session whenever new imaging conditions are met. A good approach is to train the model on videos that are heavily misclassified; for example, if the ambient lighting of the recording has changed, the classifier may perform less accurately, and those recordings should be used for retraining. For the experiments reported here, the model was retrained four times until it was robust enough that retraining was no longer required. The user can create training datasets by means of a Matlab script (movie_Labeler.m) which serves as an interface to manually classify each frame from a video of a bee trial. Usually, for a video of 300 frames, this process requires no more than 30 seconds (see instructions included in the script). All the information about the architecture of the model can be retrieved from the PER_CNN.ipynb file or after loading the model into Matlab.

### Hardware controller code

This code is written in C using the Arduino IDE. The provided code (PER_device.ino) controls the movement of the revolver by activating the stepper motor, and the feeder by activating the servo motor. All the pin assignments must match the wiring of the Arduino (Fig. S2). The code defines how long the feeder will deliver the consumable stimuli to the bee, or how to move the feeder if the tactile stimulus of the antennae should be excluded. The code for controlling the variable flux valve for the mechanical stimulus is in the file PER_AIR_Flux.ino. The valve was connected to a PWM pin on a separate Arduino which receives the input signal from the USB NI DAQ port.

### Experimental protocol

1. Collect bees and anesthetize them one by one on ice.
2. Using small tweezers, grasp the thorax and insert the neck of the bee into the slot on the bee mount (Fig. S7b).
3. With the head holder (Fig. S7a), push the head of the bee forward until the holder is fully overlapping with the top of the mount, then retract it slightly (this releases tension on the neck of the bee). The posterior edge of the head holder should line up with the back of the bee mount.
4. Place a small piece of sponge behind the body such that the bee is confined and lightly restrained (Fig. S7c).
5. Insert the bee mount on a post of the revolver, facing outward (Fig. S1b).
6. Repeat steps 2-5 for all subjects.
7. Allow the bees to fully wake up, then check for the innate PER by stimulating the antennae with sucrose. Only keep bees with a strong PER.
8. Feed all bees with 3 μl of a 50% w/v sucrose solution and place the revolver on the device. Let rest for approximately one hour.
9. Open the PER_Protocol.m file and initialize the USB NI DAQ port; next, power on the arduino and all the components.
10. Initialize the camera, and then proceed manually to reorient each bee holder such that the head of the bee is properly oriented. The head must be pointing straight to the feeder. The level of positioning precision here depends very much on the quality and mechanical tolerances of your components.
11. Mount the cover and turn on the air flow, which is necessary for the stimuli and the vacuum.
12. Adjust the feeder position and fill the feeding vessels (for these, we used p1000 pipette tips) with the desired solutions. Insert the stick (a toothpick trimmed at the proper length) for the antenna touch and soak the sponge with sucrose solution. Test that the stick passes over the head of the bee close enough to touch the antennae, and that the feeders are within reach of the bees without risk of collision.
13. Set the protocol parameters and center the heads of each bee as guided by the software.
14. Run the experiment, keeping the feeder refilled with sucrose solution as it is depleted by the bees.
15. At the end the bees can be released or maintained on the mount overnight (which requires them to be fed ad libitum following the experiment).
16. Clean the surfaces of the machine with ethanol, especially the two feeder pipette tips.

## Statistical analysis

The machine learning classifier provides for each frame a probability of proboscis extension which replaces the classical binary yes/no categorization. Statistical analysis can now be performed on this continuous variable testing its dependence on the stimuli and its temporal dynamics. Both are analyzed with a two-way repeated measures ANOVA, with CS+/- group as the between-subject variable and trials the within-subject variable. This provides total group effects and the interaction between groups and trials. Simple between-subject group effects are calculated for each trial and corrected by controlling the false discovery rate (FDR) via the Benjamini & Hochberg method. Learning- and experience-induced changes manifest themselves in simple within-subject effects in the individual groups.

## Acknowledgements

We are grateful to the Fondazione Edmund Mach for providing apicultural support. E.T. acknowledges financial support from CIMeC.

## Author Contributions

E.T. conceived the project. E.T. and H.S. developed the methods and conducted the experiments. E.T. provided private funding and instrumentation. A.H. provided funding and instrumentation. All authors contributed to the preparation of the manuscript.

## Data availability statement

With the exception of the neural network and the video recordings, all data supporting the published results (including experimental data, software for experimental control and analysis, technical drawings of mechanical parts, and electrical circuits) are publically available in the following repository: https://github.com/NeurophysicsTrento/Automized-PER

## References

1. Sasakura H, Mori I. 2013 Behavioral plasticity, learning, and memory in C. elegans. Curr Opin Neurobiol 23, 92–99. (doi:10.1016/j.conb.2012.09.005)

2. Takeda K. 1961 Classical conditioned response in the honey bee. Journal of Insect Physiology 6, 168–179. (doi:10.1016/0022-1910(61)90060-9)

3. Bitterman ME, Menzel R, Fietz A, Schäfer S. 1983 Classical conditioning of proboscis extension in honeybees (Apis mellifera). J Comp Psychol 97, 107–119.

4. Giurfa M, Sandoz J-C. 2012 Invertebrate learning and memory: Fifty years of olfactory conditioning of the proboscis extension response in honeybees. Learn Mem 19, 54–66. (doi:10.1101/lm.024711.111)

5. Matsumoto Y, Menzel R, Sandoz J-C, Giurfa M. 2012 Revisiting olfactory classical conditioning of the proboscis extension response in honey bees: A step toward standardized procedures. Journal of Neuroscience Methods 211, 159–167. (doi:10.1016/j.jneumeth.2012.08.018)

6. Frost EH, Shutler D, Hillier NK. 2012 The proboscis extension reflex to evaluate learning and memory in honeybees (Apis mellifera): some caveats. Naturwissenschaften 99, 677–686. (doi:10.1007/s00114-012-0955-8)

7. Cholé H, Junca P, Sandoz J-C. 2015 Appetitive but not aversive olfactory conditioning modifies antennal movements in honeybees. Learn Mem 22, 604–616. (doi:10.1101/lm.038448.115)

8. Birgiolas J, Jernigan CM, Gerkin RC, Smith BH, Crook SM. 2017 SwarmSight: Real-time Tracking of Insect Antenna Movements and Proboscis Extension Reflex Using a Common Preparation and Conventional Hardware. J Vis Exp, 56803. (doi:10.3791/56803)

9. Mathis A, Mamidanna P, Cury KM, Abe T, Murthy VN, Mathis MW, Bethge M. 2018 DeepLabCut: markerless pose estimation of user-defined body parts with deep learning. Nat Neurosci 21, 1281–1289. (doi:10.1038/s41593-018-0209-y)

10. Gascue F, Marachlian E, Azcueta M, Locatelli FF, Klappenbach M. 2022 Antennal movements can be used as behavioral readout of odor valence in honey bees. IBRO Neurosci Rep 12, 323–332. (doi:10.1016/j.ibneur.2022.04.005)

11. Shen M, Szyszka P, Deussen O, Galizia CG, Merhof D. 2015 Automated tracking and analysis of behavior in restrained insects. Journal of Neuroscience Methods 239, 194–205. (doi:10.1016/j.jneumeth.2014.10.021)

12. Kirkerud NH, Wehmann H-N, Galizia CG, Gustav D. 2013 APIS—a novel approach for conditioning honey bees. Front Behav Neurosci 7, 29. (doi:10.3389/fnbeh.2013.00029)

13. Smith BH. 1998 Analysis of Interaction in Binary Odorant Mixtures. Physiology & Behavior 65, 397–407. (doi:10.1016/S0031-9384(98)00142-5)

14. Hosler JS, Smith BH. 2000 Blocking and the detection of odor components in blends. Journal of Experimental Biology 203, 2797–2806. (doi:10.1242/jeb.203.18.2797)

15. Lee JK, Strausfeld NJ. 1990 Structure, distribution and number of surface sensilla and their receptor cells on the olfactory appendage of the male moth Manduca sexta. J Neurocytol 19, 519–538. (doi:10.1007/BF01257241)

16. Han Q, Hansson BS, Anton S. 2005 Interactions of mechanical stimuli and sex pheromone information in antennal lobe neurons of a male moth, Spodoptera littoralis. J Comp Physiol A Neuroethol Sens Neural Behav Physiol 191, 521–528. (doi:10.1007/s00359-005-0618-8)

17. Tuckman H, Patel M, Lei H. 2021 Effects of Mechanosensory Input on the Tracking of Pulsatile Odor Stimuli by Moth Antennal Lobe Neurons. Frontiers in Neuroscience 15.

18. Tiraboschi E, Leonardelli L, Segata G, Haase A. 2021 Parallel Processing of Olfactory and Mechanosensory Information in the Honey Bee Antennal Lobe. Front Physiol 12, 790453. (doi:10.3389/fphys.2021.790453)

19. Pacheco DA, Thiberge SY, Pnevmatikakis E, Murthy M. 2021 Auditory activity is diverse and widespread throughout the central brain of Drosophila. Nat Neurosci 24, 93–104. (doi:10.1038/s41593-020-00743-y)

20. Carcaud J, Otte M, Grünewald B, Haase A, Sandoz J-C, Beye M. 2023 Multisite imaging of neural activity using a genetically encoded calcium sensor in the honey bee. PLoS Biol 21, e3001984. (doi:10.1371/journal.pbio.3001984)

21. Hori S, Takeuchi H, Arikawa K, Kinoshita M, Ichikawa N, Sasaki M, Kubo T. 2006 Associative visual learning, color discrimination, and chromatic adaptation in the harnessed honeybee Apis mellifera L. J Comp Physiol A 192, 691–700. (doi:10.1007/s00359-005-0091-4)

22. Gallistel CR, Gelman I. 2000 Non-verbal numerical cognition: from reals to integers. Trends Cogn Sci 4, 59–65. (doi:10.1016/s1364-6613(99)01424-2)

23. Paoli M, Macri C, Giurfa M. 2023 A cognitive account of trace conditioning in insects. Current Opinion in Insect Science 57, 101034. (doi:10.1016/j.cois.2023.101034)

24. Chittka L, Skorupski P, Raine NE. 2009 Speed-accuracy tradeoffs in animal decision making. Trends Ecol Evol 24, 400–407. (doi:10.1016/j.tree.2009.02.010)

25. Zuo Y, Safaai H, Notaro G, Mazzoni A, Panzeri S, Diamond ME. 2015 Complementary contributions of spike timing and spike rate to perceptual decisions in rat S1 and S2 cortex. Curr Biol 25, 357–363. (doi:10.1016/j.cub.2014.11.065)

26. Paoli M, Haase A. 2018 In Vivo Two-Photon Imaging of the Olfactory System in Insects. Methods Mol Biol 1820, 179–219. (doi:10.1007/978-1-4939-8609-5_15)

27. Gold JI, Shadlen MN. 2007 The neural basis of decision making. Annu Rev Neurosci 30, 535–574. (doi:10.1146/annurev.neuro.29.051605.113038)

28. Dyer FC. 2002 The biology of the dance language. Annu Rev Entomol 47, 917–949. (doi:10.1146/annurev.ento.47.091201.145306)

29. Urlacher E, Francés B, Giurfa M, Devaud J-M. 2010 An Alarm Pheromone Modulates Appetitive Olfactory Learning in the Honeybee (Apis Mellifera). Front Behav Neurosci 4, 157. (doi:10.3389/fnbeh.2010.00157)

30. Paoli M, Albi A, Zanon M, Zanini D, Antolini R, Haase A. 2018 Neuronal Response Latencies Encode First Odor Identity Information across Subjects. J. Neurosci. 38, 9240–9251. (doi:10.1523/JNEUROSCI.0453-18.2018)

31. Smith BH, Abramson CI, Tobin TR. 1991 Conditional withholding of proboscis extension in honeybees (Apis mellifera) during discriminative punishment. J Comp Psychol 105, 345–356. (doi:10.1037/0735-7036.105.4.345)

32. Perry CJ, Barron AB, Chittka L. 2017 The frontiers of insect cognition. Current Opinion in Behavioral Sciences 16, 111–118. (doi:10.1016/j.cobeha.2017.05.011)

33. Bateson M, Desire S, Gartside SE, Wright GA. 2011 Agitated Honeybees Exhibit Pessimistic Cognitive Biases. Curr Biol 21, 1070–1073. (doi:10.1016/j.cub.2011.05.017)

34. de Bivort BL, van Swinderen B. 2016 Evidence for selective attention in the insect brain. Curr Opin Insect Sci 15, 9–15. (doi:10.1016/j.cois.2016.02.007)

35. Letzkus P, Ribi WA, Wood JT, Zhu H, Zhang S-W, Srinivasan MV. 2006 Lateralization of olfaction in the honeybee Apis mellifera. Curr Biol 16, 1471–1476. (doi:10.1016/j.cub.2006.05.060)

36. Frasnelli E, Haase A, Rigosi E, Anfora G, Rogers LJ, Vallortigara G. 2014 The Bee as a Model to Investigate Brain and Behavioural Asymmetries. Insects 5, 120–138. (doi:10.3390/insects5010120)

