## Supplementary material for "Temporal dynamics of honeybee learning and decision-making are revealed by a novel automated PER system"

**Supplementary Figure S1:** Automated setup characterization. a) Full setup showing the main elements: revolver, feeder, stepper motor and rotary counter, olfactometer, and control electronics. b) The revolver in isolation, with 12 bees mounted around the circumference. c) Feeder details including sucrose-soaked stick to touch the antennae, olfactometer output nozzle, and sucrose-soaked sponge on which the stick rests.

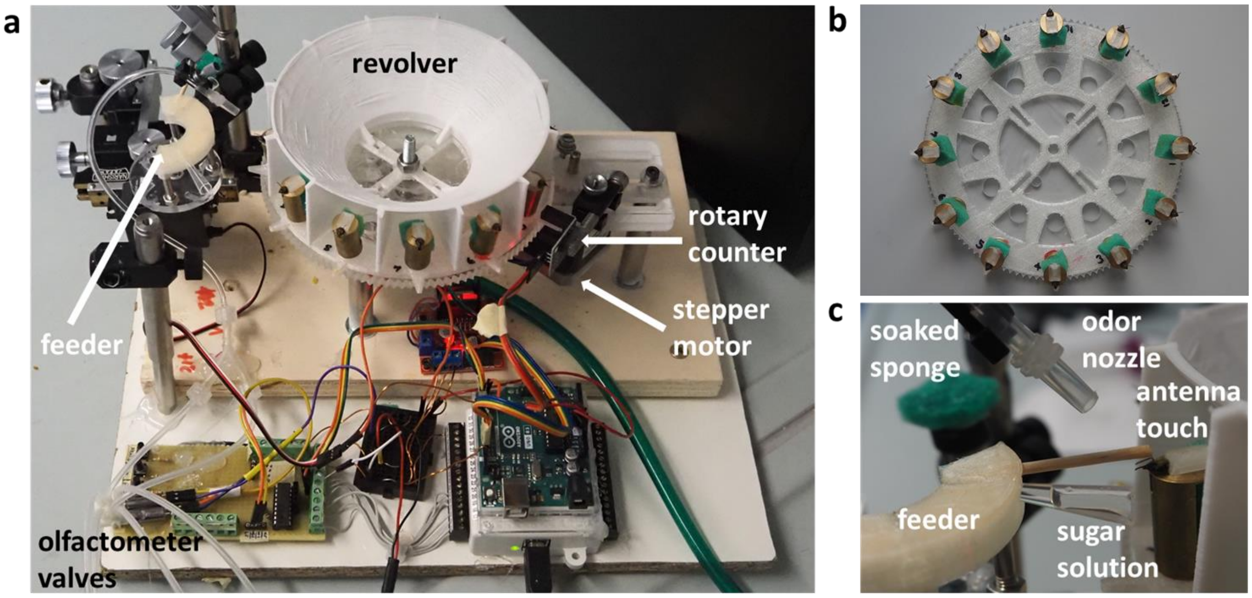


**Supplementary Figure S2:** Electronic control circuits.

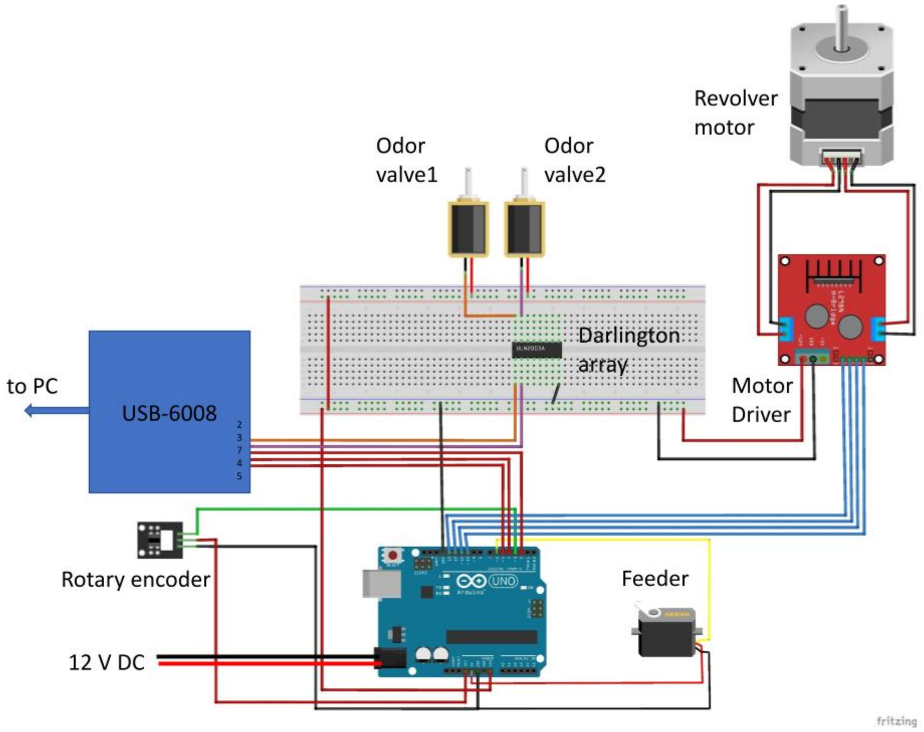

**Supplementary Table S3:** Electronic Component list.

| **component** | **function** | **producer code** |
|---|---|---|
| stepper motor driver | drive motor | L298N Dual H Bridge DC stepper Motor Drive Module |
| rotary encoder | positioning of the revolver | LM393 Photoelectric Sensor/Count Sensor |
| servo motor | feeder movements | |
| stepper motor | rotate the revolver | |
| arduino Uno | microcontroller | Arduino |
| Nidaq board | communication between the PC and the PER hardware | National Instruments usb-6008 |
| Darlington transistor array | power the actuators | ULN2003 |
| flux-controlled valve | control air flux | AP-621L-LR3-GPH, Camozzi |
| camera | recording bees | DMK27BUR0135 Imagingsource |
| lens | | 5.0-50 mm F/1.4 1/3 CS, Tamron |

**Supplementary Figure S4:** Camera positioning. The camera is positioned directly above the site of stimulation **(a)** such that the head of a given bee is centered in the left third of the field of view **(b)**, to allow for the proboscis to extend fully and remain inside the frame **(c)**.

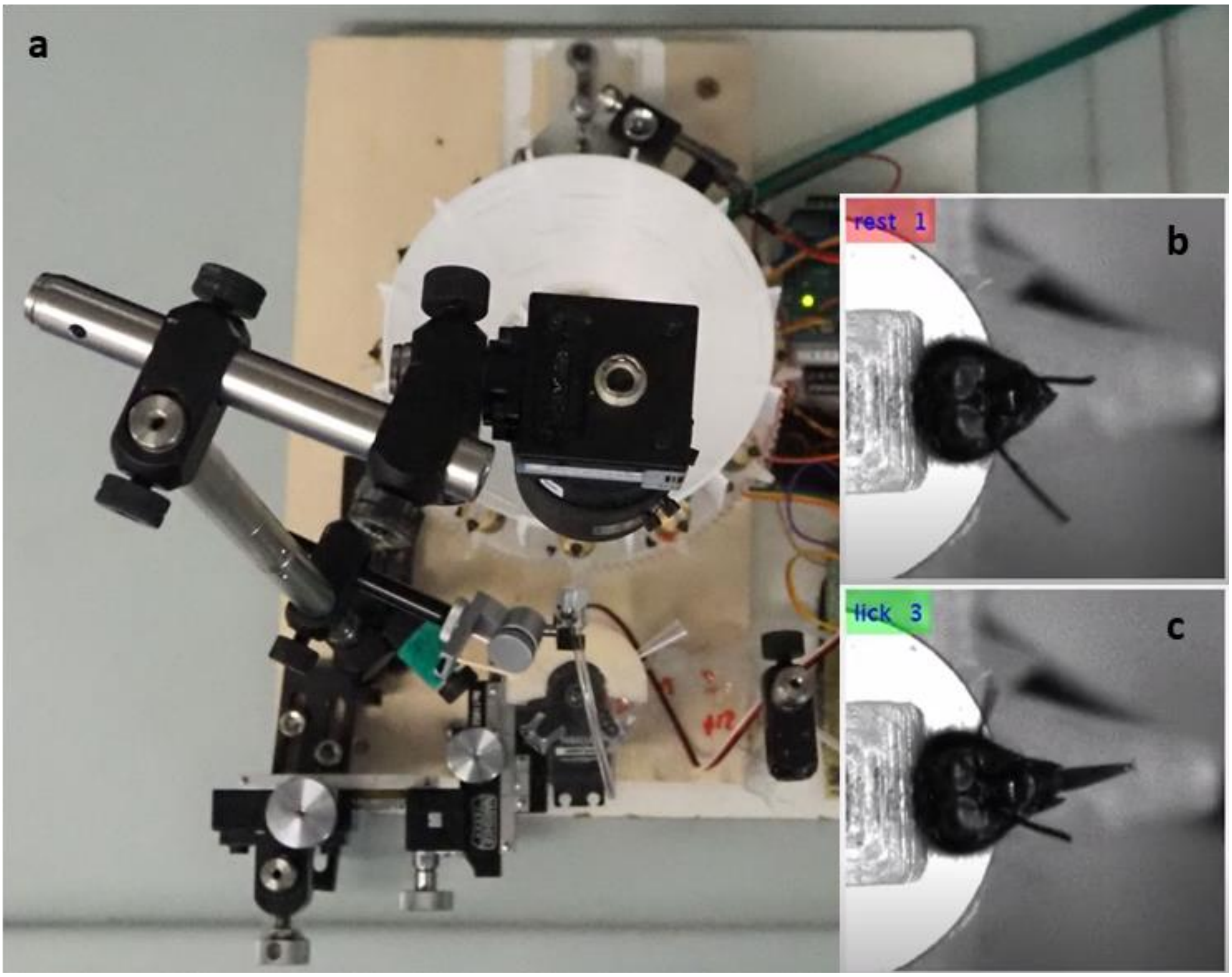

**Supplementary Figure S5:** 3D Scheme of the mechanical assembly. Left blue, Per_Mount. Left green, MOTOR_GEAR. Purple, PER_wheel. Gold, cover. Right green, UPPER_feeder. Right blue, LOWER_feeder.

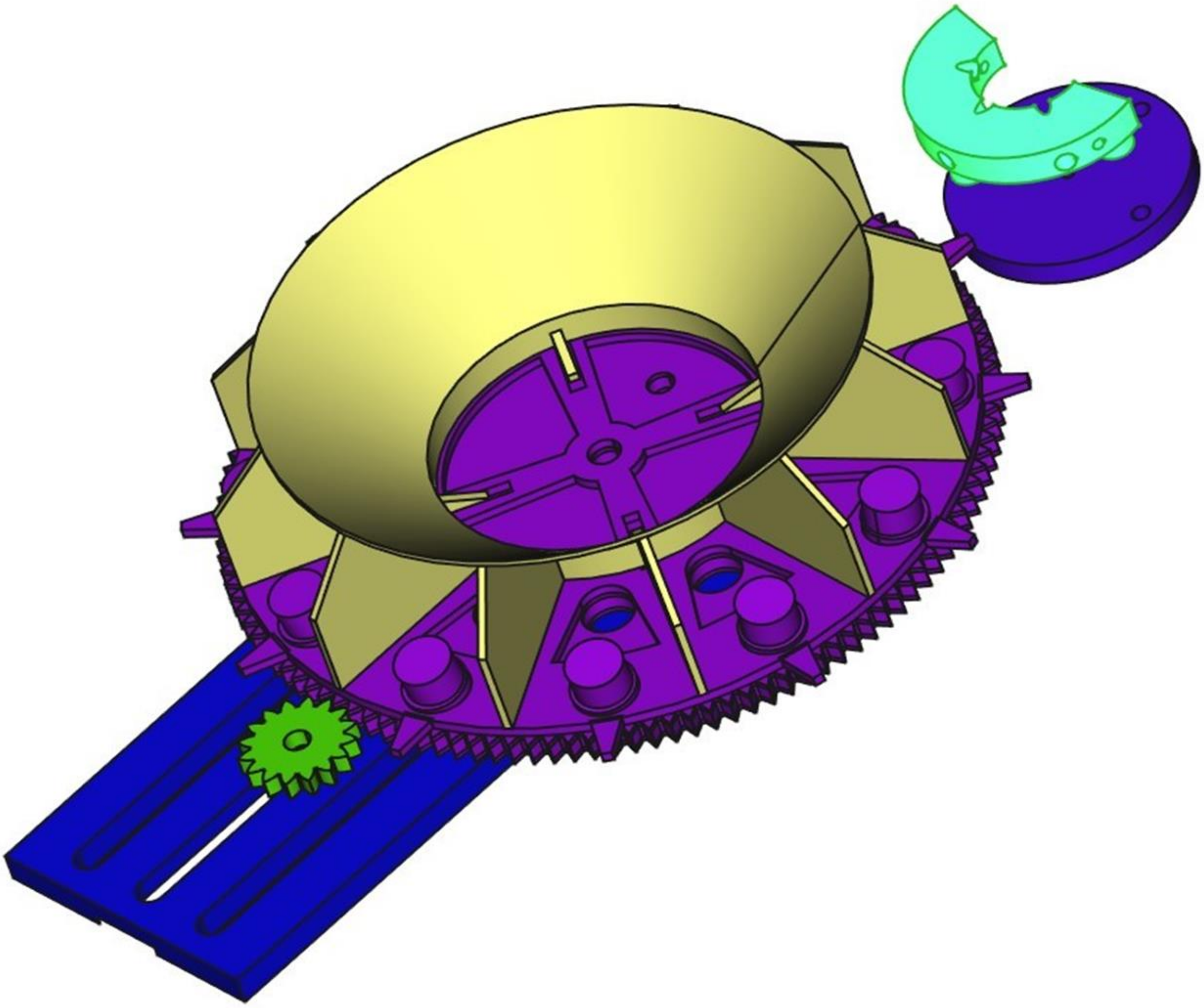

**Supplementary Table S6:** Mechanical part list.

CAD designs of all the components of the device are provided in the FCStd format, which is a FreeCAD document. They can be opened and modified with the open source FreeCAD software. Each part can be 3D printed and some of them machined with a desktop CNC.

- PER_assembly_drawing: this is the full design containing all the mechanical components. For each part, STL files are provided as well as the full assembly for rapid 3D visualization.
- per _revolver: this drawing contains multiple parts: a motor gear to be mounted on the stepper motor's shaft, the revolver, and the cover used to isolate the bees. The cover was 3D printed using PETG, whereas the other two parts can be 3D printed using any material.
- PER_FRAME: these elements represent the frame onto which the stepper motor and the revolver will be mounted. The shaft can be an M5 bolt that runs through the base and the underbase. 2 bearings are placed in the slots in the center of these 2 last pieces. The base and the underbase can be joined by means of 3 bolts, after drilling and threading of correspondent holes in the base.
- feeder: there are two pieces, the servo_ mount which is fixed onto the servo motor shaft and the feeder_mount which is connected to the servo mount with three rods of 4 mm in diameter. The distance between the two parts should be adjusted to fit the whole structure.

- head_holder: this is the piece we use for fixing in place the head of the bee. It should be 3D printed using a soft polymer like TPU.
- bee_MOUNT: this is the piece where bees are fixed for the experiment.

**Supplementary Figure S7:** Bee mounting. The head holder **(a)** slides into the notch at the top of the bee mount **(b)** to secure the bee in place for handling and experimentation **(c)**.

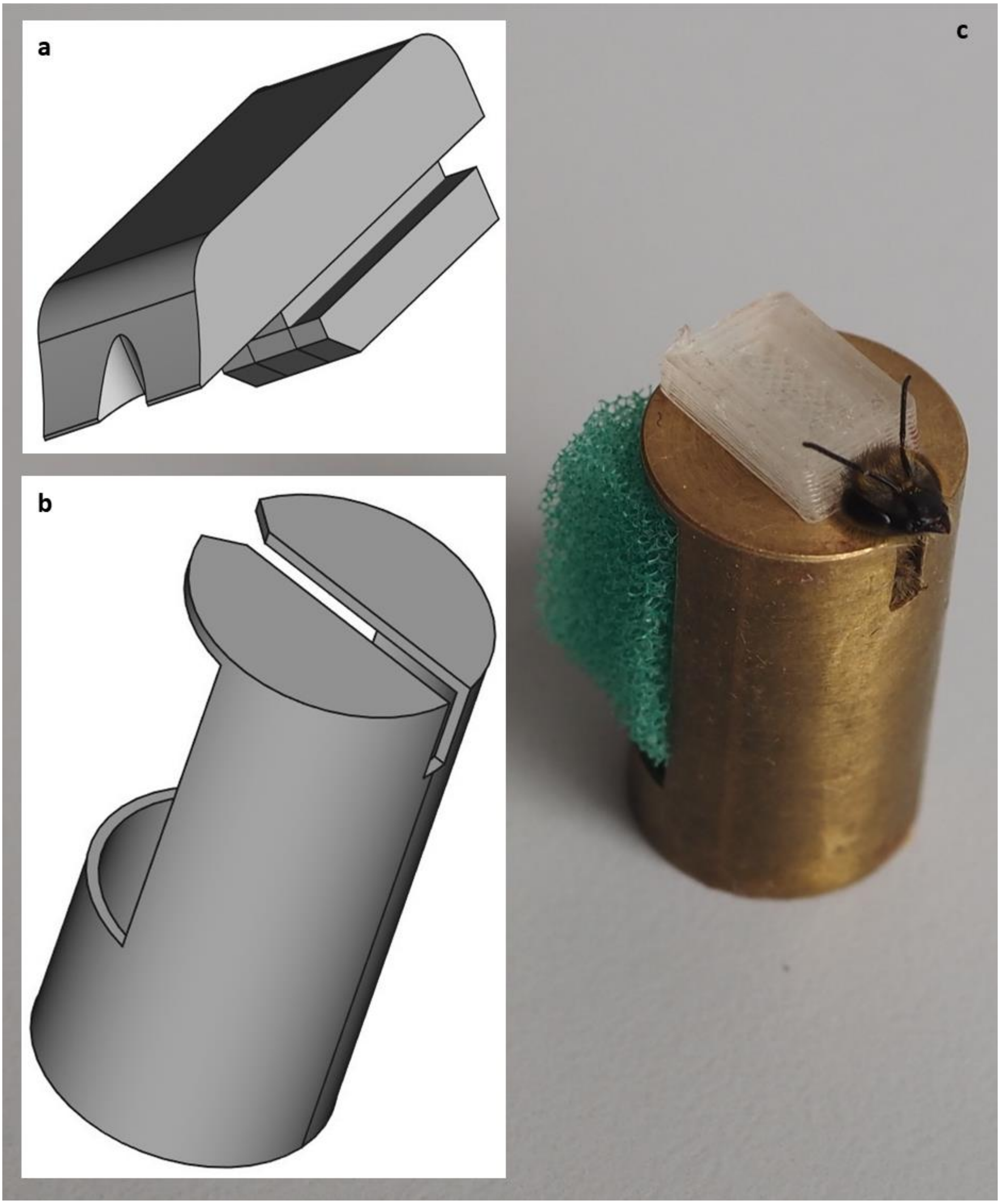

**Supplementary Video S8:** Experimental sequence.

A video showing the camera's recording of a single trial. After 2 seconds the CS+ (an odor, in this case 1-hexanol) is applied, and after 5 seconds the US is added, by first touching the antennae with a sucrose-soaked stick and then providing sucrose solution from the feeder. The inset shows the time in seconds and its colors indicates the result of the machine learning classification of whether the proboscis is extended or not (lick/rest).

**Supplementary Figure S9:** Statistical results for the trial-by-trial analysis of between-subject effects.

**Odor conditioning**

| | **Trial 1** | **Trial 3** | **Trial 3** | **Trial 4** | **Trial 5** |
|---|---|---|---|---|---|
| ***F*(1,130)** | 0.0152 | 4.93 | 33.9 | 43.4 | 60.3 |
| ***p*** | 0.9 | 0.035 | 7.3E-08 | 2.6E-09 | 1.1E-11 |

**Low-flux CS+**

| | **Trial 1** | **Trial 3** | **Trial 3** | **Trial 4** | **Trial 5** | **Trial 6** | **Trial 7** | **Trial 8** |
|---|---|---|---|---|---|---|---|---|
| ***F*(1,130)** | 0.257 | 2.51 | 1.84 | 3.86 | 2.54 | 4.41 | 5.3 | 6.64 |
| ***p*** | 0.61 | 0.13 | 1.90E-01 | 7.00E-02 | 1.30E-01 | 0.056 | 0.038 | 0.023 |
| | **Trial 9** | **Trial 10** | **Trial 11** | **Trial 12** | **Trial 13** | **Trial 14** | **Trial 15** | **Trial 16** |
| ***F*(1,130)** | 6.17 | 8.69 | 11.2 | 8.44 | 14.9 | 14.9 | 15.7 | 6.84 |
| ***p*** | 0.027 | 0.012 | 0.0049 | 0.012 | 0.0012 | 0.0012 | 0.0012 | 0.023 |

**High-flux CS+**

| | **Trial 1** | **Trial 3** | **Trial 3** | **Trial 4** | **Trial 5** | **Trial 6** | **Trial 7** | **Trial 8** |
|---|---|---|---|---|---|---|---|---|
| ***F*(1,130)** | 2.36 | 2.01 | 8.89 | 15.5 | 13 | 7.91 | 4.89 | 5.1 |
| ***p*** | 0.24 | 0.24 | 2.00E-02 | 2.60E-03 | 4.10E-03 | 0.02 | 0.068 | 0.068 |
| | **Trial 9** | **Trial 10** | **Trial 11** | **Trial 12** | **Trial 13** | **Trial 14** | **Trial 15** | **Trial 16** |
| ***F*(1,130)** | 7.84 | 0.324 | 2.09 | 1.97 | 0.314 | 0.897 | 0.351 | 1.58 |
| ***p*** | 0.02 | 0.58 | 0.24 | 0.24 | 0.58 | 0.43 | 0.58 | 0.28 |
